# Non-invasive hormone assessment of Australian Merino Rams (*Ovis aries*): A pilot investigation of cortisol, testosterone and heat stress

**DOI:** 10.1101/2024.06.12.598752

**Authors:** Dylan Fox, Benn Wilson, Edward Narayan

## Abstract

Non-invasive hormone assessment is growing in interest as producers and livestock researchers seek new methods to assess animal welfare. Conventional matrices used for hormone assessment include blood serum, saliva, urine and faeces, typically involving invasive human-animal interaction, which is known to initiate an acute stress response and thus confound studies assessing cortisol. As such, these matrices are most appropriate as short-term, point measures as they reflect circulating concentrations at the level of the minute and hour. Alternatively, non-invasive hair and wool assessment offer long-term, historic reflections of hormone concentration at the scale of weeks and months – and are not limited by sampling stress – thus making wool an appropriate tissue for hormone analysis. This pilot study quantified cortisol and testosterone concentrations of ram fleece and determine if there is a significant difference between segments of the sample staple, and whether there is a correlation between hormones. Cortisol is a glucocorticoid produced within the adrenal glands and secreted in anticipation of or in response to a stressor. Testosterone is an androgen mainly synthesised within the testes of males and responsible for several critical functions including regulation of muscle growth, libido and spermatogenesis. In our study, 70 topknot wool samples were collected from rams on a commercial stud property in Dirranbandi, Queensland, Australia. Of these animals, 12 samples were selected at random to undergo cortisol and testosterone quantification. In the laboratory, a single, intact staple was isolated from the total sample, divided into 10 mm segments and prepared for their respective (cortisol or testosterone) immunoassays. No significant difference (p>0.05) was found between wool segments for either cortisol or testosterone, however, statistical differences (p<0.05) were found between individuals for both hormones. A strong correlation (R2=0.9173) was found between wool cortisol and testosterone concentrations, which was a first-time discovery in rams. Furthermore, climate loggers (n=6) were installed in proximity to the rams to collect daily maximum temperature (°C) and relative humidity (RH%) data to calculate the temperature-humidity index (THI) as an indicator of potential heat stress. Under this method, rams were deemed ‘comfortable’ at a THI<72; under ‘mild’ stress at a THI of between 72 and 78; ‘moderate’ stress between 79 and 80; and ‘severe’ stress at a THI of >81. Mean weekly THI peaked in late summer (February), remained high in early autumn (March), decreased throughout the remainder of autumn and the duration of winter before increasing slightly as temperatures rose in early spring (September). Over the trial, 90.36% or 4,706 h were marked by ‘comfortable’ conditions, 8.95% or 466 h by ‘mild’ stress, 0.60% or 31 h by ‘moderate’, and 0.10% or 5 h by ‘severe’ stress. It was determined that THI is most valuable when combined with other methods of measuring heat stress, including non-invasive wool hormone assessment. Whilst most of the findings in this study were previously confirmed by other studies, the strong correlation between wool cortisol and testosterone concentrations appears to be a first. In summary, this study reveals the major future possibilities for non-invasive wool hormone assessment and the possible applications of combining this with temperature-humidity index scores to provide further insight into heat stress within the context of production animal agriculture.

## Introduction

Cortisol is a steroid hormone in the class of glucocorticoids, colloquially referred to as the ‘stress hormone’ for its role in the stress response (Davenport et al. 2006; Russell et al. 2012). Cortisol actively circulates the body via blood and is found in numerous bodily tissues, including saliva, excreta (urine and faeces), hair and wool. Wool hormone analysis is a non-invasive method useful for retrospective studies due to the delay in hormone incorporation into wool.

Cortisol analysis provides an indication of activation of the HPA axis which mediates the stress response, with support from the SAM axis. The stress response is a natural reaction to the perception of a threat and initially attempts to maintain homeostasis despite the stressor (Russell et al. 2012; Salaberger et al. 2016). Acute stress is a short-acting activation of the stress response, whilst chronic activation results from prolonged stress. Both can have significant effects on production livestock (Sejian et al. 2018). Testosterone is critical for the proper development of male reproductive system and as such can also be assessed.

Instances of heat stress can also be quantified and qualified without the need for hormonal assessment. The temperature-humidity index (THI) provides and indication of the severity of heat stress using climatic data.

An acute stress response is a short-lived reaction to a perceived threat, and includes routine animal husbandry tasks, including shearing and drafting (Yardimci et al. 2013; Stubjoen et al. 2015). Early work by Matteri et al. (1984) states rams subjected to temporarily (3 h) restraint stress reported lower than basal levels of LH. As gonadotrophins stimulate the testes to secrete testosterone, this suggests even short-lived, acute stressors can impact animal productivity. Furthermore, Doey et al. (1976) found 6 h of wet stress disrupted the LH surge that accompanies ovulation in ewes. These findings support the notion that acute stress can disrupt the reproductive systems of both rams and ewe.

In instances of prolonged or repeat stress, chronic activation of the stress response may occur. This process is often cumulative, whereby several occasions of HPA activation compound, resulting in the accumulation of cortisol within the bloodstream, and over time, other bodily tissues (Russell et al. 2012). Repeated exposure to a single stressor can result in a habitual HPA axis response (Dhabhar et al. 1997). Tilbrook et al. (2000) also noted marked declines in gonadotrophin secretion during prolonged stress. Interestingly, not all stress responses are equally severe. Figueiredo et al. (2003) found predation stress resulted in a long-term, consistent release of glucocorticoid. Herman et al. (2016) warns that researchers should hesitate to define chronic stress solely by the activity (or inactivity) of the HPA axis as several systems, including the autonomic nervous and immunes systems, form part of the stress response. In summary, the stress response system is multifaceted and highly dependent on the specifics of the animal’s context, including their nutrition status, environmental conditions and production stage.

Early research on growth rate and feed utilisation efficiency found ram lambs outperformed their castrated counterparts and attributed the difference to testosterone (Walker, 1950; Schanbacher et al. 1980). An androgen and the primary male sex hormone, testosterone is critical for the normal maturation of male animals as it promotes several critical developments, including protein synthesis, contributing to the greater size of males; increased bone density; and for promotion of proper penis, scrotal and testes development (Hamilton et al. 2016). It is also necessary for correct spermiogenesis, which is critically important in siring male.

In seasonal breeders, including the ram, reproductive axis activation occurs in relation to day length (Thibault, et al. 2001). Declining daylight stimulates oestrus behaviour in ewes and elevated testosterone production in males (Perkins and Roselli, 2007). Testosterone is fundamentally important for libido and continual sexual activity by rams. Thwaites (1982) recognised that sexual activity increases alongside testosterone levels. The opposite was also proven true with castrated ram lambs exhibiting little sexual behaviour in adulthood (Parrott, 1978). Tilbrook et al. (1993) found similar results in a study where prepubescent rams were administered a GnRH agonist which resulted in decreases in circulating testosterone, likely the result of overstimulation and eventual desensitisation of the anterior pituitary to GnRH and a decline in FSH and LH secretion (Vickery et al. 1986). Interestingly, Knight (1973) found no significant correlation between circulating blood testosterone and ram libido, suggesting factors other than androgen levels affect ram sex drive.

Heat stress is a fundamental issue across many production systems due to Australia’s extreme climate. The vulnerability of production livestock to heat stress varies by breed, age, nutritional and production status (Das et al. 2016). For instance, heritage breeds are often far more resilient to local climatic conditions compared to exotic breeds due to many generations of genetic adaption (Rojas-Downing et al. 2017). Heat stress is known to affect all aspects of production including growth (Bohmanova et al. 2007), milk yield (Das et al. 2016) and reproductive performance (Oduwole et al. 2021). As predicted by current climate research, instances of heat stress are likely to become both more common and intense (Sejian et al. 2018). As such, heat stress research is gaining significant attention. In their heat stress review, Sejian and colleagues (2018) emphasises the need for new biological markers for measuring heat stress. The authors explain via their model that as heat stress often compounds other stressors – including limited access to feed and water – one or more of the production functions are sacrificed to provide the energy necessary to combat the stressor (Sejian et al. 2018).

Developed in the 1990s, the temperature-humidity index (THI) is a calculation of a single value using temperature and humidity data to estimate the levels of environmental heat stress animals are subjected to. Different versions of the formula are used for different production species due to differences in physiology and associated cooling mechanisms (Bohmanova et al. 2007). Humidity data is critical as the amount of water vapor in the air has a direct effect on an animal’s ability to cool itself as the rate of evaporative loss is diminished. THI is considered a reliable predictor of the severity of heat stress; however, two significant drawbacks exist (Sejian et al. 2018). THI calculations exclude both wind speed and solar radiation measures, which influence the ‘feels like’ or apparent temperature. To overcome these limitations, Gaughan et al. (2008) developed the ‘Heat load index’.

The aim of this study was to critically evaluate the utility of non-invasive wool hormone assessment within the context of heat stress. This investigation occurred in isolation of conventional, invasive methods uses to assess hormone concentrations. The study objective are 1) evaluate whether there is a statistically significant difference between wool cortisol and testosterone across the length of the staple; and 2) assess the merit of temperature-humidity index (THI) values. It is hypothesised that no statistically significant difference will be found within the staple for either hormone, and THI will provide an indication of the amount and severity of heat stress experienced by the rams.

## Methods

### Ethics approval

Ethics approval was approved by the University of Queensland (2022/AE000424).

### Field work

#### Animals and husbandry

All 70 rams used in this study were owned by Wilgunya Merino Stud located in Dirranbandi, Queensland, 4486 (−28.856123236319117, 148.4624339024027). At the time of our visit (14th October 2022), all rams were housed in a separate paddock with ad libitum access to natural grasslands and fresh drinking water. Property managers used ATVs and working dogs to herd the rams into the race for sample collection.

#### Wool collection

Rams were run through the race where an identifying tag number was noted on both a resealable bag and a strip of paper, which was placed within the bag. Handheld electric clippers were used to recover a sample of topknot fleece which was placed within its own labelled bag. 70 samples of ram fleece were collected. All samples were collected by the farm support worker to reduce risk of animal injury by unexperienced personnel. Upon return to the University of Queensland, Gatton Campus, 4343 (−27.55331668592056, 152.3344294537852), all fleece ram fleece samples were placed into a large vacuum-sealable bag and placed within a Wayco freezer at - 20oC.

### Laboratory analysis

#### Wool preparation

Methodology used adapted from Sawyer et al. (2019). Before use, the vacuum-sealed bag was reinflated and allowed to reach room temperature before samples were catalogued. Of the 70 samples, a random selection of 12 were selected for analysis in this study. From each of these 12 samples, an intact, clean, unstretched staple was identified and cut out. This piece was deemed the representative subsample for that individual moving forward. All other samples were returned to the freezer.

Using a ruler marked with 1 millimetre (mm) increments, each subsample was cut into 10 mm pieces and assigned to a labelled weighing boat (see Appendix A). Samples were labelled as ‘animal ID number’ and a letter, ranging from A to G. For instance, 70C represented the third piece from ram #70. Sample ‘A’ was furthest to the scalp and ‘G’ the closest. The number of 10 mm subsamples varied between rams due to differences in total staple length. The weight of all samples – minus their boat – were recorded using a digital balance (Ohaus PioneerTM) accurate to three decimal places. In an effort to avoid human error, the balance was connected to a laptop via an RS-232 cable to allow for the direct transmission of weight data into Microsoft Excel.

#### Washing procedure

Early findings by Davenport et al. (2006) using hair and subsequent studies using wool recommended the use of isopropanol as a washing agent to remove external contamination from the sample surface (Sawyer et al. 2019). As such, all wool subsamples were submerged in 3 millilitres (mL) of 100 % isopropanol for five minutes, drained and placed within a glass desiccator until dry. Samples typically dried within 48 hours of washing and were then diced finely using forceps and surgical scissors and added to an Eppendorf tube with 1 mL of 100% ethanol for hormone extraction. Samples were briefly vortexed for at least one second to ensure maximum ethanol penetration of finely cut wool. Tubes were labelled and placed within a refrigerator.

#### Cortisol assay

Cortisol concentration was determined using the DetectX® Cortisol Immunoassay kit, manufactured by Arbor Assays. Briefly, a pipette was used to extract 500 μL of each sample into new, labelled Eppendorf tubes. Tubes were placed in a fume hood with the caps open to dry overnight. Remaining methanol sample was held in the refrigerator in case samples needed to be rerun. Once dry, samples were reconstituted with 25 μL of 100% methanol and 475 μL of ELISA assay buffer which was prepared by diluting assay buffer concentration provided in the DetectX® Cortisol Immunoassay kit with distilled water in the ratio 1:5.

NuncTM 96-well plates were coated with 50 μL of cortisol antibody solution and incubated for at least 12 hours. Standards were prepared serially using 200 μL of standard stock and 200 μL of assay buffer.

Four non-specific binding wells were used and included 75 μL of assay buffer was used and 50 μL of assay buffer to the two maximum binding wells (labelled ‘0’ on the plate map; see Appendix B). 25 μL of DetectX® Cortisol Conjugate was added to every well along with 25 μL of DetectX ® Cortisol Antibody, which was omitted from the NSB wells.

Once loaded, the plate was covered using sealing film, labelled with the assay type, student details and time, and set atop the plate shaker at 900 rpm for 1 hour. Next, the plate was washed X times using an automatic plate washer. Once washed, the plate was gently tapped dry using paper towel. 100 μL of TMB Substrate was added to each well, the plate resealed and left to incubate at room temperature for 30 minutes. 50 μL of stop solution was added to each well. The plate was read at 450 nm using a microplate reader. Cortisol concentrations were exported as a CSV. Data was transformed from absorbance values (AU cm-1) into cortisol concentrations provided in nanograms per gram (ng/g). Cortisol standards are run in duplicate from 400, 200, 100, 50, 25, 12.5, 6.25, 3.12, 1.56. Data obtained from transformed into the standard curve which is used to determine the accuracy of our result, post-assay (see Appendix C and Appendix D).

#### Testosterone assay

Testosterone concentration was determined using the R156/7 enzyme within an immunoassay. A 96-well NuncTM Maxisorp plate was coated the day prior to the assay (see Appendix E). 25 μL of antibody stock was added to 6225 μL of coating buffer, at a working dilution of 1:25,000. The first column were reserved as non-specific binding wells and were coated with coating buffer an no antibody. 50 μL antibody was added per well using a multi-pipette. The plate was gently tapped on the table to maximise antibody coverage, covered and left for at least 12 hours at 4°C.

Standards, including zeros and NSB, were prepared the following morning. Standard values were run in duplicate from 100, 50, 25, 12.5, 6.25, 3.12, 1.56, 0.78, 0.39 pg/well. Standard working stock was diluted serially two-fold using 200 μL stock and 200 μL EIA buffer. This was repeated for all remaining standards. Samples were diluted in EIA buffer. 15 μL of testosterone HRP was added to 5985 μL of EIA buffer to create the working dilution.

An automatic plate washer was used to wash the plate four times with wash solution. Paper towel was used to gently dry any remaining wash solution from the plate after washing. Using the plate map as a guide, 50 μL of standard, control and sample were added per well. 50 μL of diluted testosterone HRP was added to all wells that contained standard, control or sample. Within 10 minutes of beginning, the loaded plate was covered and left to incubate at room temperature for two hours. Immediately after this time, the plate was washed four times with was solution using the automatic plate washer. The plate was briefly inverted to remove excess solution and carefully dried with paper towel.

50 μL of TMB substrate was added to wells that contained standard, control or sample. The plate was covered with adhesive film and left to shake for 30 minutes. The plate had now developed colour. 50 μL of stop solution was added to wells that contained standard, control or sample. Plate was inserted into plate reader at read at 450 nm. Absorbance values were exported as a CSV, imported into Microsoft Excel and transformed to testosterone concentration (ng/g).

### Temperature-humidity index

Six climate loggers (HOBO U23 Pro v2) were set up within a ∼200 mm radius of the property’s farmstead on February 6^th^, 2022 by previous research students. The location was an estimated 1 kilometre from the paddock where the rams resided. Climate loggers recorded temperature and relative humidity (minimum, maximum and average for both) hourly for 217 days, before their batteries depleted. Four of six climate loggers were retrieved from the property on October 14th 2022; two were not located.

Logger data was downloaded as four CSV files, imported into Microsoft Excel and analysed. Of the four devices, loggers 8 and 12 were selected at random. Data recorded outside the ranges of February 7th, 2022 – 11th September, 2022 were incomplete and thus discarded.

Temperature-humidity index (THI) was calculated using the formula;

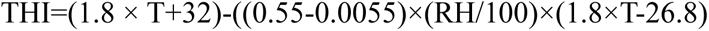

as provided by Tucker et al. 2008. This paper also that a THI value <72 falls within the ‘comfort’ range for small ruminants (i.e. sheep and goats), THI = 72-78 suggests ‘mild’ stress, THI =79-80 ‘moderate’ stress and THI = ≥ 81 ‘severe’ stress.

THI scores were calculated for the climate logger 8 and 12 dataset and averaged to find the overall mean maximum THI for 217 days and then the weekly means. Note: THI calculations used both maximum temperature (°C) relative humidity (%) data provided from loggers 8 and 12. This decision was intentional to identify all hours of potential heat stress. Use of minimum or average data would fail to recognise temperature extremes, and thus provide less accurate data.

### Statistical analysis

#### Hormone study

Cortisol and testosterone absorbance data (AU cm^-1^) was provided by the plate reader as a CSV which was transformed into both cortisol and testosterone concentration (ng/g) in Microsoft Excel. First analysis included descriptive statistics including mean, standard deviation, standard error. A few outlier values were found; these are provided below for transparency. All such values were removed from all calculations.

#### Cortisol samples removed included

Errors resulted in the removal of sample 5E, 15D, 41C, 57D.

Two values were provided for 29A, despite only having one position on the plate map (likely technician error).

Two values were provided for 70A, despite only have one position on the plate map. Also, sample 70D had a reading of 96.78 ng/g, over 12-times higher than the next value for that ram. As such, ram #70 was struck from cortisol analysis.

#### Testosterone samples removed included

Errors resulted in the removal of 5E and 5D.

#### Climate data

Raw climate logger data was imported into Microsoft Excel as a CSV file and transformed into THI values using basic descriptive statistics, including mean, range, and maximum. Initial graphs were made in Excel.

All data were checked for homogeneity of variances and log-transformed prior to analysis. Statistical analysis included non-parametric ANOVA to compare mean levels of WTC and WCC between sub-samples and Ram ID. Spearman correlation was done for comparison between WT and WCC. Line graph and bar chart were used to plot climate and THI data.

## Results

### Wool cortisol

No statistically significant (p>0.05) difference was found between an individual’s samples; however, a statistically significant (p<0.05) difference was found between individuals. WCC varied widely between individuals, ranging from 0.25 ng/g (Ram 38C) to 68.7 ng/g (Ram 29C). Mean WCC for each sheep is presented in **Figure 1**.

**Figure 1.**
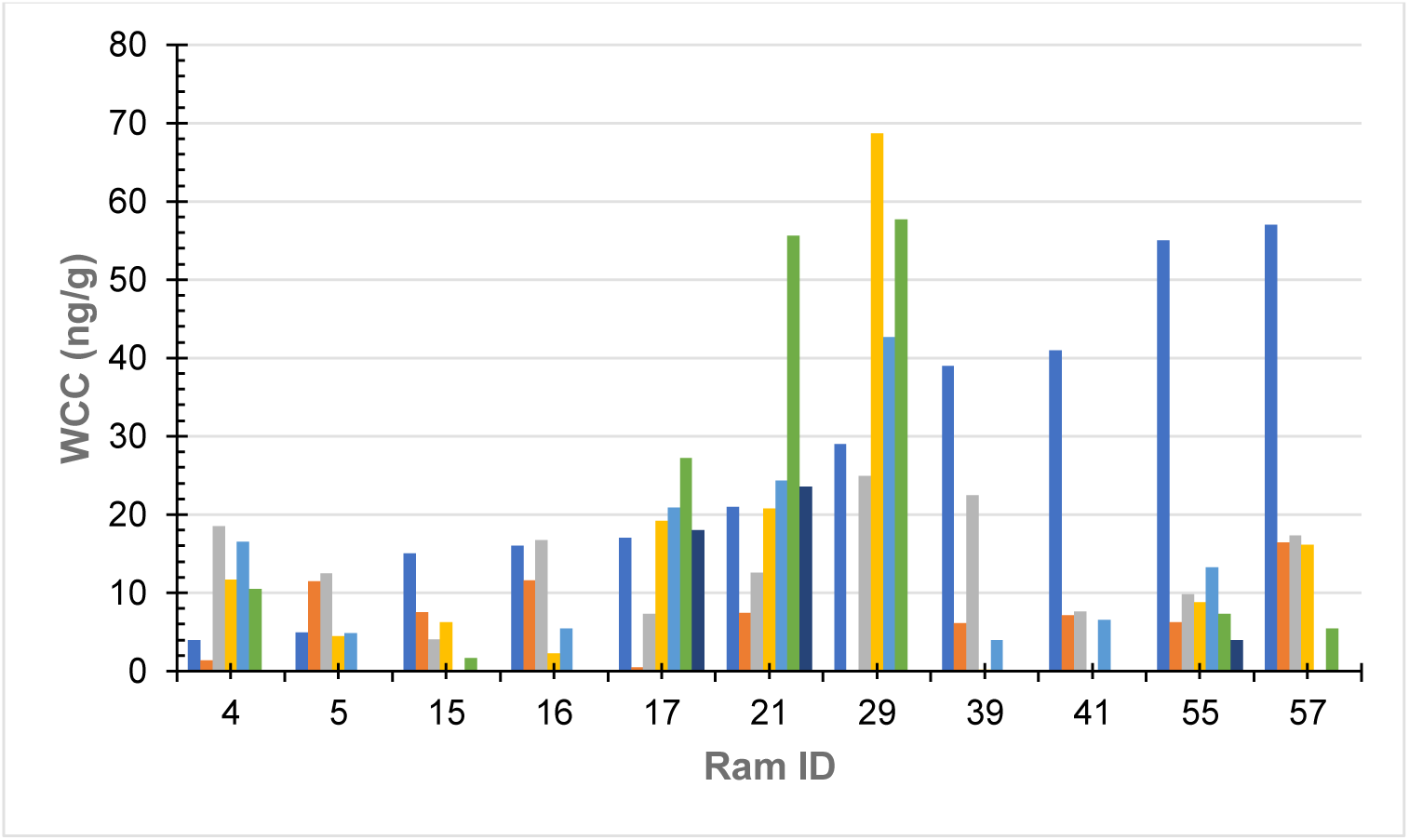
Wool cortisol concentration by segment.

### Wool testosterone

No statistically significant (p>0.05) difference was found between an individual’s samples; however, a statistically significant (p<0.05) difference was found between individuals. WTC varied wildly between individuals, ranging from 0.08 ng/g (Ram 55E) to 11.6 ng/g (Ram 29C). Mean WTC for each sheep is presented in **Figure 2**.

**Figure 2.**
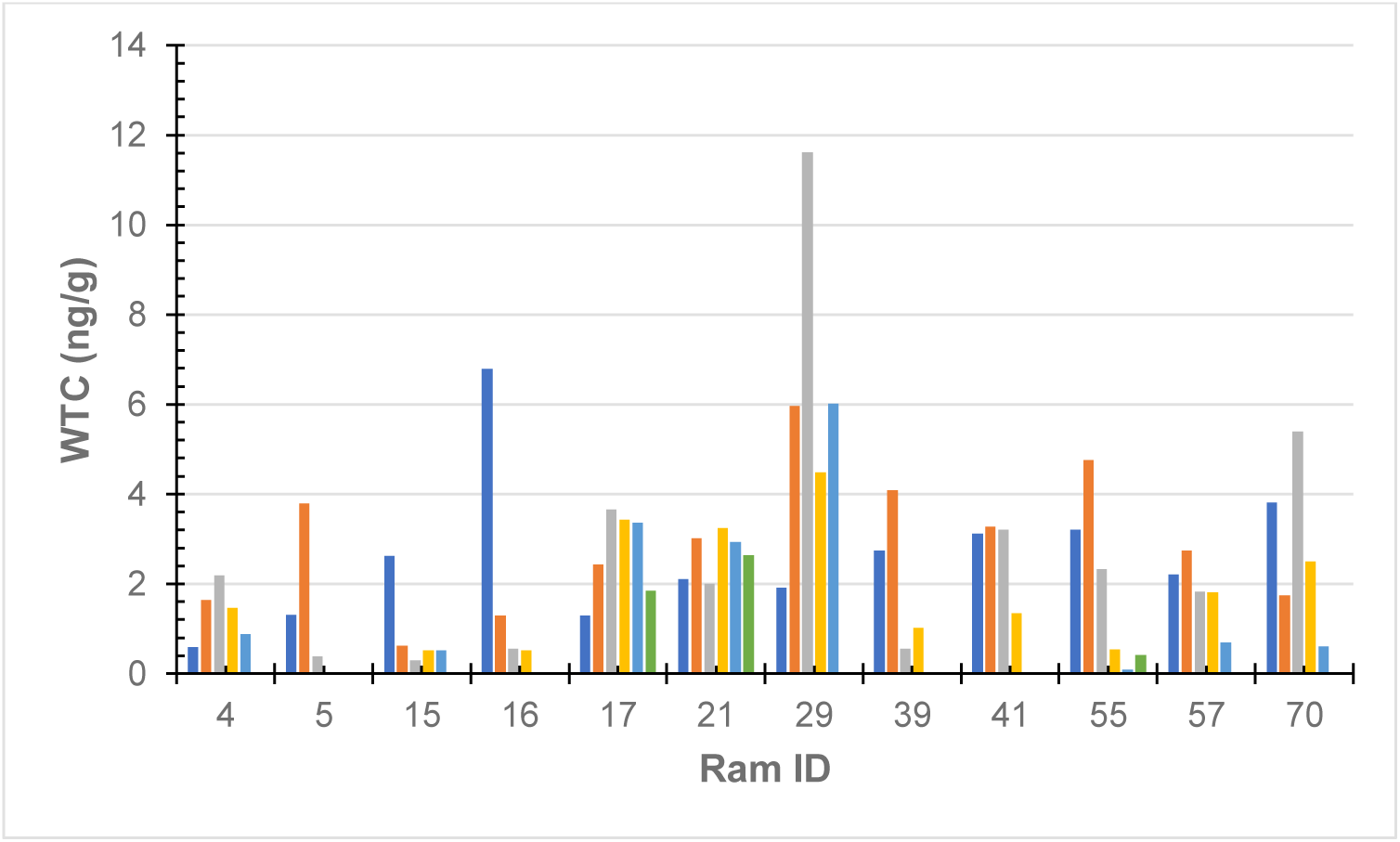
Wool testosterone concentration by segment.

### Wool cortisol and testosterone

In Figure 3 below, X-axis represents ram (n=11) identification number and y-axis the hormone concentration, measured in ng/g. Error bars depict standard error (SE) of each data set, which measures variability around the mean.

**Figure 3.**
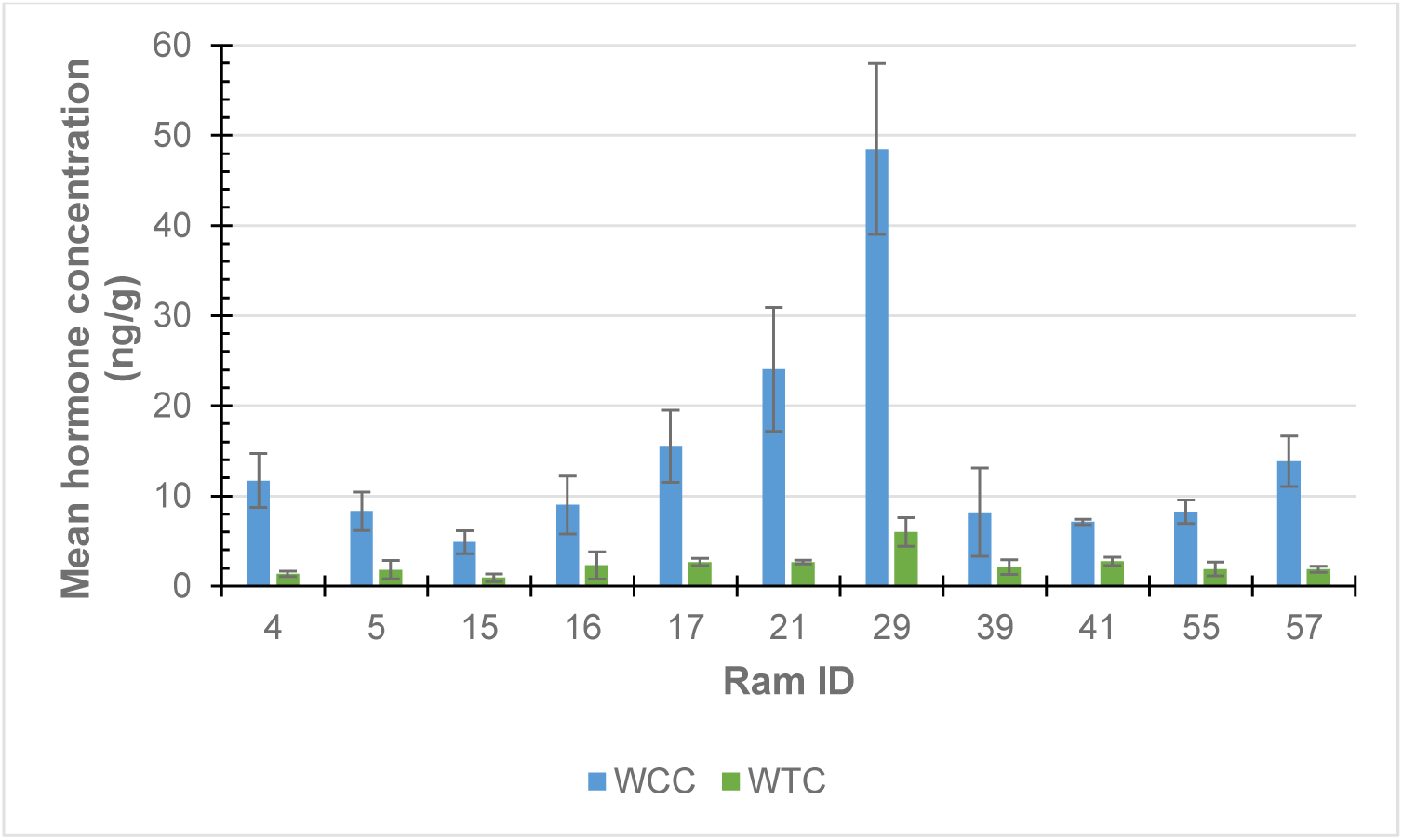
Ram wool mean cortisol (WCC; blue) and testosterone (WTC; green) concentrations.

Scatter plot represents mean WCC (n=11) and mean (n=11) for rams #4, 5, 15, 16, 17, 21, 29, 39, 41, 55 and 57. WTC data for ram #70 was omitted as the WCC data point was deemed an outlier. The linear regression intercept line is determined by the equation y = 0.2245x, suggesting WTC increases by 0.2245 ng/g per ng/g of WTC, on average. The R^2^ of 0.9173 suggests a strong, positive correlation between both variables (Figure 4).

**Figure 4.**
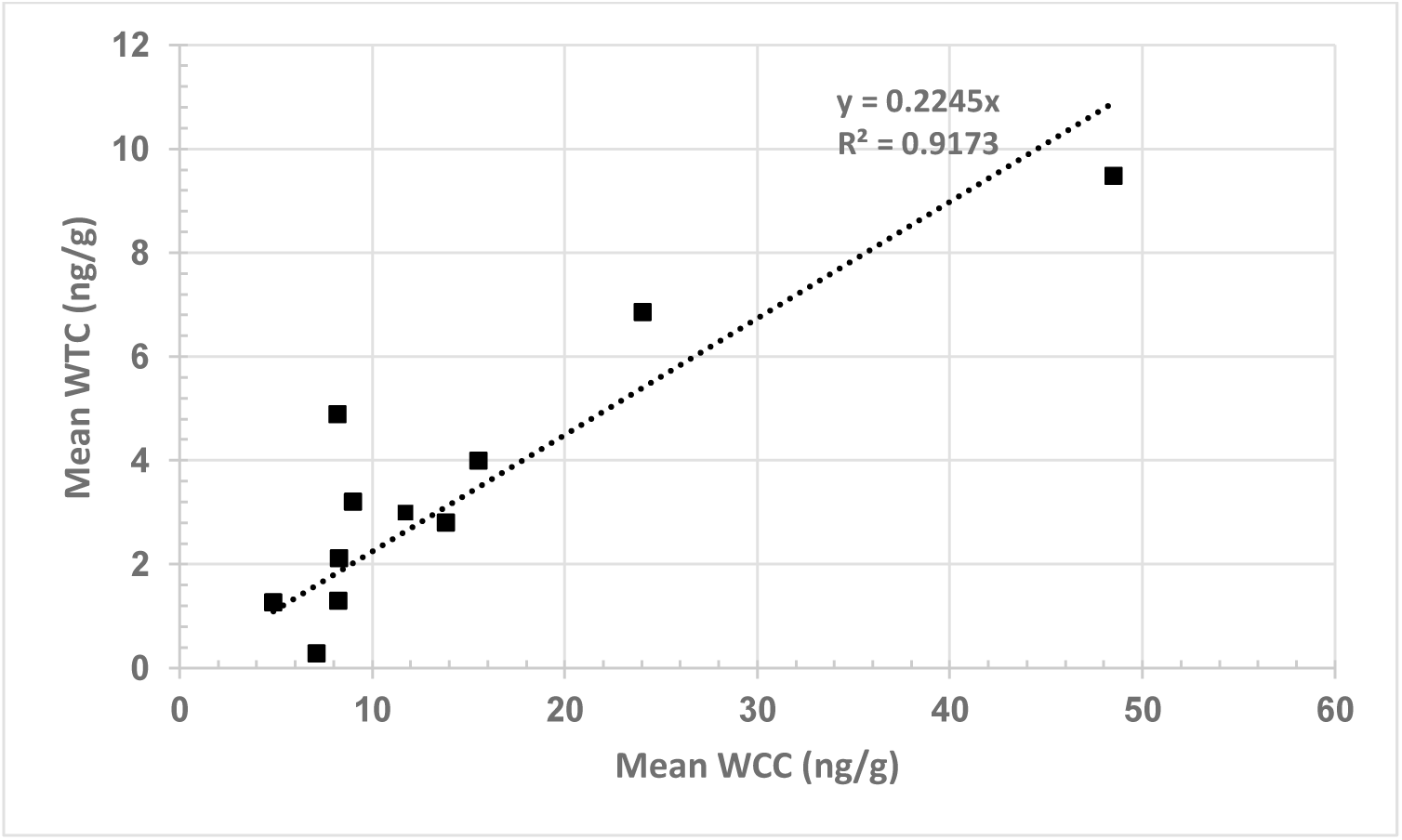
Relationship between wool cortisol concentration (WCC; ng/g) and wool testosterone concentration (WTC; ng/g).

### Temperature-humidity index

Line graph (Figure 5) shows mean (n=2) weekly THI fluctuations for the duration of the trial. The x-axis represents months and is subdivided into weeks data was available. The y-axis represents the mean maximum THI score. Note the y-axis is truncated (beginning at a THI of 60) to increase resolution of data variation over time. Data was calculated by first finding the mean weekly maximum THI for Logger 8 and 12, and then calculating the mean. Data presented ranges from 7^th^ February-11^th^ September 2022.

**Figure 5.**
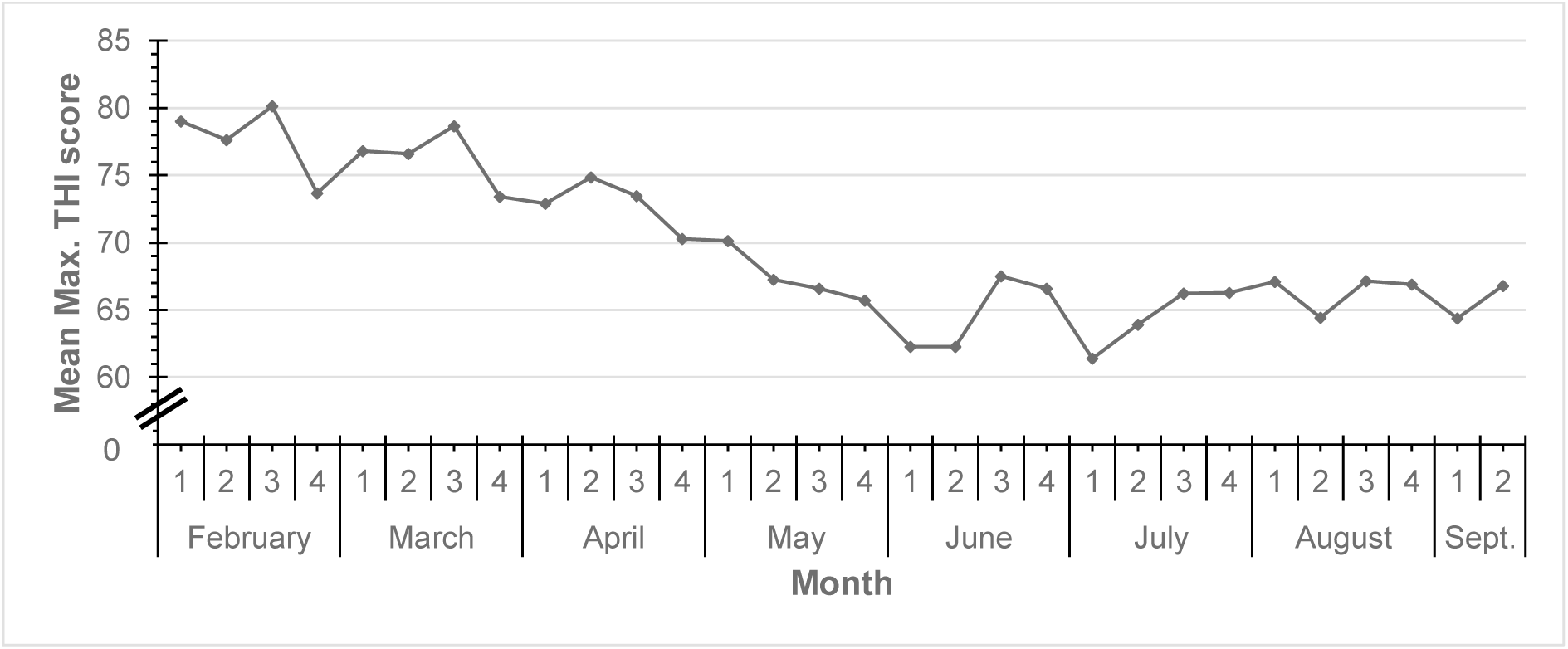
Weekly variation in mean maximum temperature-humidity index (THI) scores.

Figure 6 represents cumulative hours at each level, with Comfortable (THI<71) in green, Mild (72-78) in yellow, Moderate (78-80) in orange and Severe (≥81) in red. The sum total of hours for each month is provided on the right of each month. Data presented ranges from 7^th^ February-11^th^ September 2022.

**Figure 6.**
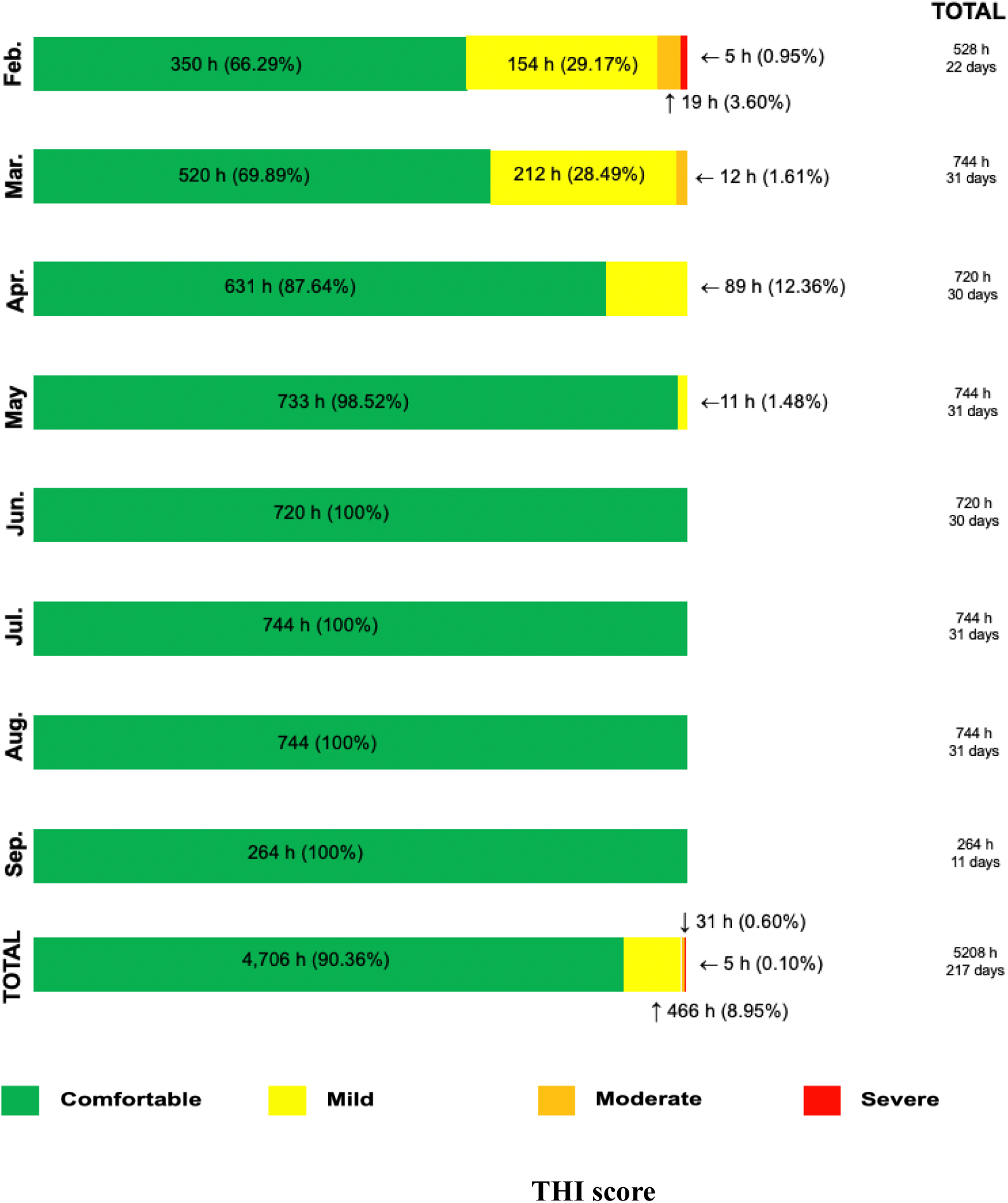
Aggregate time (hours) at each level of THI predicted heat stress.

## Discussion

### Wool hormone assessment

This study assessed the variation in WCC and WTC along the length of the staple. Hormone concentrations of 10 mm staple segments were determined by hormone-specific enzyme immunoassays as per an adapted hair analysis protocol. Mean ram WCC findings are presented in Figure 1, mean ram WTC in Figure 2 and combined in Figure 3. No statistical difference (p>0.05) in either wool cortisol nor testosterone were found along the length of the staple. These findings are concurrent with Hantzopoulou et al. (2022) who reported no statistically significant difference in the wool cortisol concentration when subsampling Merino ewe topknot fleece. Similarly, Davenport et al. (2006) found no significant difference (p>0.05) in cortisol between distal and proximal portions of monkey hair. These results are replicated in a human hair study by Yang et al. (1998) which found no significant difference in hair cortisol at three different lengths. Caution, however, must be applied during inter-species comparison as several biological differences are at play. For instance, the use of human hair products is known to alter the hormone profile of hair. Nonetheless, it demonstrates that – to date – the assessment of hair segments is yet to yield a statistically significant difference, at least under mild stressors.

Similar to our colleagues, the present study did find a statistically significant difference (p<0.05) in WCC and WTC between animals, demonstrating that the hormonal activity of individual rams varies within the flock. Hormones are chemical secretions that facilitate communication between bodily systems. Cortisol is the primary hormone of the mammalian stress response and is released by the adrenal glands, where it enters the bloodstream. Whilst the exact mechanism of incorporation is disputed, strong evidence supports the passive diffusion of hormones into the hair shaft via blood (Russell et al. 2012). As such, wool hormone concentration varies animal to animal as a direct reflection of the level of HPA axis activity.

Our hypothesis, which suggested that there is no statistically significant difference in both wool cortisol and testosterone along the length of the staple, was proven correct. Additionally, we also confirmed that wool hormone concentrations vary significantly between individuals. As the rams were cohabitating under similar environmental and much effort was taken to eliminate potential stressors, it is reasonable to expect similar hormone results, as each animal was experiencing comparable levels of HPA axis activity. Had the rams been subjected to extreme heat stress, the data would reflect this.

An early iteration of this study attempted to assess the validity of using hair as a long-term historical record of ram heat stress. Insufficient climate data and resource constraints resulted in the termination of that element of the project. However, it is noted that the findings of the wool study would have likely proven true. As there was no significant difference across the length of the staple (∼6 cm long), it is unlikely segmented cortisol values would relate to normal climate. It is hypothetically possible that a severe, chronic stressors, such as a week-long heat wave, water deprivation or extreme pestilence, may lead to a strong spike in systematic cortisol and would later be reflected in analysed wool. After an extensive literature search, it is believed that this research is the first investigation of ram wool testosterone.

### Wool cortisol and testosterone correlation

Within many mammalian animal breeding populations, males actively compete against rival males in a dominance hierarchy for access to females. It is often the case that testosterone levels are related to the number of dominance dispute they have won, and thus their position in the dominance hierarchy (Medill et al. (2023). Cortisol levels, however, vary across species with examples of dominant males exhibiting low and higher blood glucocorticoid concentrations.

A positive correlation (R2 = 0.91) was found between WCC and WTC, which appears to be the first documented case of such a relationship in rams (Figure 4). A study by Medill et al. (2023) found a non-significant, positive correlation (p>0.05; R2 =0.43) in hair cortisol and testosterone in feral horses. Bachelors (males of breeding age yet to win access to their own mare) were hypothesised to have lost more dominance disputes and thus have elevated cortisol levels (Medill et al. 2023). Whilst such a strong correlation is interesting – particularly in social animals, such as rams – it is difficult to confidently identify the exact reasoning for such a relationship from a single wool sample. Moreover, as wool samples were collected in October 2022, it is highly likely that seasonal changes that accompany the breeding season (February to June) are reflected in the samples and contribute to intra-sample hormone variability.

Future iterations of this study should first determine if seasonal hormone variability is reflected in wool, as they are in conventional blood samples. Secondly, in-field observations should be conducted to identify dominant rams within the flock and later through repeat sampling and data analysis determine how wool cortisol and testosterone change alongside shifts in dominance hierarchy positioning.

### Temperature-humidity index (THI) and heat stress

The temperature-humidity index incorporates ambient temperature and relative humidity into an equation to determine the severity of heat stress experienced by livestock. The formula reported by Tucker et al. (2008) was used and divided THI scores into four categories, including ‘comfortable’, ‘mild’, ‘moderate’, and ‘severe’ heat stress. Weekly mean maximum THI for the duration of the trial is presented in Figure 5 and unsurprisingly reflects the seasonal variation in temperature and relative humidity for the Dirranbandi field site. Instances of heat stress are typically more common during summer and early autumn, which occurs from December to February in the Southern Hemisphere. The THI calculations prepared in this study reflect this trend, with February and March recording the highest mean weekly THI scores for the entire data set, respectively. Whilst Figure 5 shows a clear seasonal trend in THI score, it fails to identify the key instances of severe heat stress, as the graph reflects mean weekly THI, rather than maximum weekly THI. Greater resolution is provided in Figure 8 where the aggregate time at each level of stress is provided for each month.

Similar to the previous figure, Figure 6 clearly shows progressive temperature declines as depicted by the growing proportion of ‘comfortable’ conditions over time. Overall, based on these calculations, rams experienced comfortable conditions for over 90 percent of the time and just under 9 percent of time at a moderate level. Trial rams were at or near comfort level for over 99 percent of the 4,706 h data was collected. This result is of significant importance as instances of prolonged heat stress can have severe and lasting impacts on animal performance, ranging from reduced ability to thermoregulate, reduced average daily gain, stunted wool growth rate and poor spermatogenesis, which can have lasting repercussions for future flock size (Sejian et al. 2018). The latter is of particular concern for the trial rams as they were part of a Merino ram stud that serviced a growing ewe flock. Hamilton et al. (2015) observed decreases in sperm motility and significant increases in major and minor spermatozoa defects in rams that had undergone simulated heat stress. Critically, semen quality remained impaired for 47 days, or roughly equivalent to one sperm cycle. Whilst this is far outside the scope of the present study, a future investigation could determine whether THI score is a reliable indicator of semen quality, within the context of heat stress. If true, the calculation of THI scores may enable producers to forecast the effects of severe heat stress on the reproductive fitness of their ram flock.

This reveals that trial rams experienced 5 hours of severe heat stress during February, equivalent to 0.95 percent of the month and 0.1 percent of the total trial time. As data was only collected between late summer and early spring, it is highly likely that several severe heat stress events were not captured by climate loggers. The limited collection of spring and summer data was a major limitation of this study. Loggers were established near the field site in early February 2022 and collected hourly data until mid-September 2022. As such, this captured the entirety of autumn and winter – a time unlikely to have any lasting heat stress events. It is noted that heavy rains and local flooding limited travel to and from the field site during late 2021 and as such loggers were established as soon as feasible. Future investigations of THI should focus on identifying instances of heat stress during the warmer months.

## Conclusion

Wool provides a suitable matrix for assessment of ram cortisol and testosterone. No statistically significant differences were found along the length of the staple for either hormone. These findings were confirmed by literature. As hypothesised, a statistically significant difference was found between individuals. Measuring wool cortisol and testosterone provide an indication of the activity of the HPA and male HPG axis. Moreover, a strong correlation was found between wool cortisol and testosterone concentrations. After extensive literature search, it is believed this study is the first to find such a correlation in wool. Finally, THI proved a valuable indicator of potential severe heat stress. Limited access to spring and summer data proved a limitation of this study. Further studies should assess the relationship between wool cortisol concentration and THI to determine how both techniques can better predict heat stress in Australian merino rams.

## Acknowledgements

We would like to acknowledge the contributions made by The Australian Wool Education Trust (AWET) for the scholarship and partial funding for this project. This research was conducted as part of Masters of Animal Science research project for Dylan Fox and supervised by Dr Edward Narayan with support provided by Professor John Gaughan during manuscript draft stages. Members of the Stress Lab including Harsh Pahuja and Georgia-Constantina Hanztopoulou provided peer support with lab work. We would like to acknowledge Max Wilson of Wilgunya Merino Stud for the collaboration.

## References

Ansari-Renani, H., & Hynd, P. (2001). Cortisol-induced follicle shutdown is related to staple strength in Merino sheep. Livestock Production Science, 69(3), 279–289.

Ashley, N., Barboza, P., Macbeth, B., Janz, D., Cattet, M., Booth, R., & Wasser, S. (2011). Glucocorticosteroid concentrations in feces and hair of captive caribou and reindeer following adrenocorticotropic hormone challenge. General and Comparative Endocrinology, 172(3), 382–391.

Barnes, A., Beatty, D., Taylor, E., Stockman, C., Maloney, S., & McCarthy, M. (2004). Physiology of heat stress in cattle and sheep. Meat and Livestock Australia, 209, 1–36.

Bohmanova, J., Misztal, I., & Cole, J. B. (2007). Temperature-humidity indices as indicators of milk production losses due to heat stress. Journal of dairy science, 90(4), 1947–1956.

Bouwknecht, J. A., Olivier, B., & Paylor, R. E. (2007). The stress-induced hyperthermia paradigm as a physiological animal model for anxiety: a review of pharmacological and genetic studies in the mouse. Neuroscience & Biobehavioral Reviews, 31(1), 41–59.

Cannon, W. B. (1915). Bodily changes in pain, hunger, fear, and rage. D. Appleton and company.

Cannon, W. B. (1932). Homeostasis. The wisdom of the body. Norton, Newyork.

Cavigelli, S. A. (1999). Behavioural patterns associated with faecal cortisol levels in free-ranging female ring-tailed lemurs, Lemur catta. Animal behaviour, 57(4), 935–944.

Chan, J., Sauvé, B., Tokmakejian, S., Koren, G., & Van Uum, S. (2014). Measurement of cortisol and testosterone in hair of obese and non-obese human subjects. Experimental and Clinical Endocrinology & Diabetes, 122(06), 356–362.

Cobb, T., Hantzopoulou, G.-C., & Narayan, E. (2023). Relationship between wool cortisol, wool quality indices of Australian Merino Rams and climatic variables in Tasmania. Frontiers in Animal Science, 4, 1234343.

Collier, R. J., Gebremedhin, K., Macko, A. R., & Roy, K. S. (2012). Genes involved in the thermal tolerance of livestock. Environmental stress and amelioration in livestock production, 379–410.

Cone, E. J. (1996). Mechanisms of drug incorporation into hair. Therapeutic drug monitoring, 18(4), 438–443. https://oce.ovid.com/article/00007691-199608000-00022/PDF

Creel, S., Wildt, D. E., & Monfort, S. L. (1993). Aggression, reproduction, and androgens in wild dwarf mongooses: a test of the challenge hypothesis. The American Naturalist, 141(5), 816–825.

Curtis, A., Scharf, B., Eichen, P., & Spiers, D. (2017). Relationships between ambient conditions, thermal status, and feed intake of cattle during summer heat stress with access to shade. Journal of thermal biology, 63, 104–111.

Davenport, M. D., Tiefenbacher, S., Lutz, C. K., Novak, M. A., & Meyer, J. S. (2006). Analysis of endogenous cortisol concentrations in the hair of rhesus macaques. General and Comparative Endocrinology, 147(3), 255–261.

Dettmer, A. M., Novak, M. A., & Meyer, J. S. (2023). Are hair cortisol levels dependent on hair growth rate? A pilot study in rhesus macaques. General and Comparative Endocrinology, 340, 114308.

Dhabhar, F. S., McEwen, B. S., & Spencer, R. L. (1997). Adaptation to prolonged or repeated stress–comparison between rat strains showing intrinsic differences in reactivity to acute stress. Neuroendocrinology, 65(5), 360–368.

Doney, J., Gunn, R., Smith, W., & Carr, W. (1976). Effects of pre-mating environmental stress, ACTH, cortisone acetate or metyrapone on oestrus and ovulation in sheep. The Journal of Agricultural Science, 87(1), 127–132.

Figueiredo, H. F., Bodie, B. L., Tauchi, M., Dolgas, C. M., & Herman, J. P. (2003). Stress integration after acute and chronic predator stress: differential activation of central stress circuitry and sensitization of the hypothalamo-pituitary-adrenocortical axis. Endocrinology, 144(12), 5249–5258.

Fürtbauer, I., Solman, C., & Fry, A. (2019). Sheep wool cortisol as a retrospective measure of long-term HPA axis activity and its links to body mass. Domestic Animal Endocrinology, 68, 39–46.

Gaughan, J., Mader, T. L., Holt, S., & Lisle, A. (2008). A new heat load index for feedlot cattle. Journal of Animal Science, 86(1), 226–234.

Godoy, L. D., Rossignoli, M. T., Delfino-Pereira, P., Garcia-Cairasco, N., & de Lima Umeoka, E. H. (2018). A comprehensive overview on stress neurobiology: basic concepts and clinical implications. Frontiers in behavioral neuroscience, 12, 127.

Goldstein, D. S. (1987). Stress-induced activation of the sympathetic nervous system. Bailliere’s clinical endocrinology and metabolism, 1(2), 253–278.

Goldstein, D. S. (2006). Adrenaline and the inner world: an introduction to scientific integrative medicine. JHU Press.

Hamilton, T. R. d. S., Mendes, C. M., Castro, L. S. d., Assis, P. M. d., Siqueira, A. F. P., Delgado, J. d. C., Goissis, M. D., Muiño-Blanco, T., Cebrián-Pérez, J. Á., Nichi, M., Visintin, J. A., & Assumpção, M. E. O. D. Á. (2016). Evaluation of Lasting Effects of Heat Stress on Sperm Profile and Oxidative Status of Ram Semen and Epididymal Sperm. Oxid Med Cell Longev, 2016, 1687657–1687612. 10.1155/2016/1687657

Hantzopoulou, G.-C., Sawyer, G., Tilbrook, A., & Narayan, E. (2022). Intra-and inter-sample variation in wool cortisol concentrations of Australian merino lambs between twice or single shorn ewes. Frontiers in Animal Science, 3, 890914.

Haque, N., Ludri, A., Hossain, S., & Ashutosh, M. (2013). Impact on hematological parameters in young and adult Murrah buffaloes exposed to acute heat stress.

Hemsworth, P. H., Rice, M., Karlen, M. G., Calleja, L., Barnett, J. L., Nash, J., & Coleman, G. J. (2011). Human–animal interactions at abattoirs: Relationships between handling and animal stress in sheep and cattle. Applied animal behaviour science, 135(1-2), 24–33.

Herman, J. P., McKlveen, J. M., Ghosal, S., Kopp, B., Wulsin, A., Makinson, R., Scheimann, J., & Myers, B. (2016). Regulation of the hypothalamic-pituitary-adrenocortical stress response. Comprehensive physiology, 6(2), 603.

Higashiyama, Y., Nashiki, M., Narita, H., & Kawasaki, M. (2007). A brief report on effects of transfer from outdoor grazing to indoor tethering and back on urinary cortisol and behaviour in dairy cattle. Applied animal behaviour science, 102(1-2), 119–123.

Hirschenhauser, K., Möstl, E., & Kotrschal, K. (1999). Seasonal patterns of sex steroids determined from feces in different social categories of greylag geese (Anser anser). General and Comparative Endocrinology, 114(1), 67–79.

Hogan, L., Phillips, C., Horsup, A., Janssen, T., & Johnston, S. (2011). Technique for faecal marking in group-housed southern hairy-nosed wombats Lasiorhinus latifrons. Australian Zoologist, 35(3), 649–654.

Indu, S., & Pareek, A. (2015). A review: Growth and physiological adaptability of sheep to heat stress under semi–arid environment. International Journal of Emerging Trends in Science and Technology, 2(9), 3188–3198.

Kapoor, A., Schultz-Darken, N., & Ziegler, T. E. (2018). Radiolabel validation of cortisol in the hair of rhesus monkeys. Psychoneuroendocrinology, 97, 190–195.

Keckeis, K., Lepschy, M., Schöpper, H., Moser, L., Troxler, J., & Palme, R. (2012). Hair cortisol: a parameter of chronic stress? Insights from a radiometabolism study in guinea pigs. Journal of Comparative Physiology B, 182, 985–996.

Kong, L., Tang, M., Zhang, T., Wang, D., Hu, K., Lu, W., Wei, C., Liang, G., & Pu, Y. (2014). Nickel nanoparticles exposure and reproductive toxicity in healthy adult rats. International journal of molecular sciences, 15(11), 21253–21269.

Koren, L., Mokady, O., Karaskov, T., Klein, J., Koren, G., & Geffen, E. (2002). A novel method using hair for determining hormonal levels in wildlife. In (Vol. 63, pp. 403–406): Academic Press.

Mahgoub, O., Kadim, I. T., Al-Dhahab, A., Bello, R. B., Al-Amri, I. S., Ali, A. A. A., & Khalaf, S. (2010). An assessment of Omani native sheep fiber production and quality characteristics. Journal of Agricultural and Marine Sciences [JAMS], 15, 9–14.

Matteri, R., Watson, J., & Moberg, G. (1984). Stress or acute adrenocorticotrophin treatment suppresses LHRH-induced LH release in the ram. Reproduction, 72(2), 385–393.

Medill, S. A., Janz, D. M., & McLoughlin, P. D. (2023). Hair Cortisol and Testosterone Concentrations in Relation to Maturity and Breeding Status of Male Feral Horses. Animals, 13(13), 2129.

Monk, C. S., Hart, K. A., Berghaus, R. D., Norton, N. A., Moore, P. A., & Myrna, K. E. (2014). Detection of endogenous cortisol in equine tears and blood at rest and after simulated stress. Veterinary ophthalmology, 17, 53–60.

Morrow, C., Kolver, E., Verkerk, G., & Matthews, L. (2000). Urinary corticosteroids: an indicator of stress in dairy cattle. PROCEEDINGS-NEW ZEALAND SOCIETY OF ANIMAL PRODUCTION

Nejad, J. G., Lohakare, J., Son, J., Kwon, E., West, J., & Sung, K. (2014). Wool cortisol is a better indicator of stress than blood cortisol in ewes exposed to heat stress and water restriction. Animal, 8(1), 128–132.

Nejad, J. G., Sung, K., Lee, B., Peng, J., Kim, J., Oh, S., Chemere, B., & Kim, B. (2016). Effects of adding water to total mixed ration on water consumption, nutrient digestibility, wool cortisol, and blood indices in Corriedale ewes under hot and humid conditions. Journal of Animal Science, 94, 832–832.

Oduwole, O. O., Huhtaniemi, I. T., & Misrahi, M. (2021). The roles of luteinizing hormone, follicle-stimulating hormone and testosterone in spermatogenesis and folliculogenesis revisited. International journal of molecular sciences, 22(23), 12735.

Peric, T., Comin, A., Corazzin, M., Montillo, M., & Prandi, A. (2020). Comparison of AlphaLISA and RIA assays for measurement of wool cortisol concentrations. Heliyon, 6(10).

Perkins, A., & Roselli, C. E. (2007). The ram as a model for behavioral neuroendocrinology. Hormones and behavior, 52(1), 70–77.

Plotsky, P. M., Cunningham Jr, E. T., & Widmaier, E. P. (1989). Catecholaminergic modulation of corticotropin-releasing factor and adrenocorticotropin secretion. Endocrine reviews, 10(4), 437–458.

Poudel, S., Fike, J. H., & Pent, G. J. (2022). Hair Cortisol as a Measure of Chronic Stress in Ewes Grazing Either Hardwood Silvopastures or Open Pastures. Agronomy, 12(7), 1566. https://www.mdpi.com/2073-4395/12/7/1566

Pragna, P., Sejian, V., Soren, N., Bagath, M., Krishnan, G., Beena, V., Devi, P. I., & Bhatta, R. (2018). Summer season induced rhythmic alterations in metabolic activities to adapt to heat stress in three indigenous (Osmanabadi, Malabari and Salem Black) goat breeds. Biological Rhythm Research, 49(4), 551–565.

Rivier, C., & Rivest, S. (1991). Effect of stress on the activity of the hypothalamic-pituitary-gonadal axis: peripheral and central mechanisms. Biology of reproduction, 45(4), 523–532.

Russell, E., Koren, G., Rieder, M., & Van Uum, S. (2012). Hair cortisol as a biological marker of chronic stress: current status, future directions and unanswered questions. Psychoneuroendocrinology, 37(5), 589–601.

Säkkinen, H., Tornbeg, J., Goddard, P. J., Eloranta, E., Ropstad, E., & Saarela, S. (2004). The effect of blood sampling method on indicators of physiological stress in reindeer (Rangifer tarandus tarandus). Domest Anim Endocrinol, 26(2), 87–98. 10.1016/j.domaniend.2003.07.002

Salaberger, T., Millard, M., El Makarem, S., Möstl, E., Grünberger, V., Krametter-Frötscher, R., Wittek, T., & Palme, R. (2016). Influence of external factors on hair cortisol concentrations. General and Comparative Endocrinology, 233, 73–78.

Sawyer, G., Fox, D. R., & Narayan, E. (2021). Pre-and post-partum variation in wool cortisol and wool micron in Australian Merino ewe sheep (Ovis aries). PeerJ, 9, e11288.

Schanbacher, B., Crouse, J., & Ferrell, C. (1980). Testosterone influences on growth, performance, carcass characteristics and composition of young market lambs. Journal of Animal Science, 51(3), 685–691.

Schell, C. J., Young, J. K., Lonsdorf, E. V., Mateo, J. M., & Santymire, R. M. (2017). Investigation of techniques to measure cortisol and testosterone concentrations in coyote hair. Zoo Biol, 36(3), 220–225. 10.1002/zoo.21359

Sejian, V., Bhatta, R., Gaughan, J., Dunshea, F., & Lacetera, N. (2018). Adaptation of animals to heat stress. Animal, 12(s2), s431–s444.

Selye, H. (1936). A syndrome produced by diverse nocuous agents. Nature, 138(3479), 32–32.

Selye, H. (1950). Stress and the general adaptation syndrome. British medical journal, 1(4667), 1383.

Stoebel, D. P., & Moberg, G. P. (1982). Repeated acute stress during the follicular phase and luteinizing hormone surge of dairy heifers. Journal of dairy science, 65(1), 92–96.

Stradaioli, G., Peric, T., Montillo, M., Comin, A., Corazzin, M., Veronesi, M. C., & Prandi, A. (2017). Hair cortisol and testosterone concentrations and semen production of Bos taurus bulls. Italian journal of animal science, 16(4), 631–639. 10.1080/1828051X.2017.1303339

Stubsjøen, S. M., Bohlin, J., Dahl, E., Knappe-Poindecker, M., Fjeldaas, T., Lepschy, M., Palme, R., Langbein, J., & Ropstad, E. (2015). Assessment of chronic stress in sheep (part I): The use of cortisol and cortisone in hair as non-invasive biological markers. Small Ruminant Research, 132, 25–31.

Taherti, M., Issad, N. A., Khelef, D., & Mimoune, N. (2023). Reproduction Characteristics of Ouled Djellal Rams in a Semi-arid Area in Algeria.

Tan, S. Y., & Yip, A. (2018). Hans Selye (1907–1982): Founder of the stress theory. Singapore medical journal, 59(4), 170.

Thau, L., Gandhi, J., & Sharma, S. (2019). Physiology, cortisol.

Thibault, C., & Levasseur, M.-C. (2001). Reproduction in mammals and man. INRA Editions.

Tilbrook, A., Galloway, D., Williams, A., & Clarke, I. (1993). Treatment of young rams with an agonist of GnRH delays reproductive development. Hormones and behavior, 27(1), 5–28.

Tilbrook, A. J., Turner, A. I., & Clarke, I. J. (2000). Effects of stress on reproduction in non-rodent mammals: the role of glucocorticoids and sex differences. Reviews of reproduction, 5(2), 105–113.

Tucker, C. B., Rogers, A. R., & Schütz, K. E. (2008). Effect of solar radiation on dairy cattle behaviour, use of shade and body temperature in a pasture-based system. Applied animal behaviour science, 109(2-4), 141–154.

Ulrich-Lai, Y. M., Xie, W., Meij, J. T., Dolgas, C. M., Yu, L., & Herman, J. P. (2006). Limbic and HPA axis function in an animal model of chronic neuropathic pain. Physiology & Behavior, 88(1-2), 67–76.

Vázquez, R., & Orihuela, A. (2001). Effect of vaginal mucus and urine from ewes in estrus on plasma testosterone levels and weight gain of feedlot rams. Small Ruminant Research, 42(3), 171–175.

Venturelli, E., Cavalleri, A., & Secreto, G. (1995). Methods for urinary testosterone analysis. Journal of Chromatography B: Biomedical Sciences and Applications, 671(1-2), 363–380.

Verbeek, E., Colditz, I., Blache, D., & Lee, C. (2019). Chronic stress influences attentional and judgement bias and the activity of the HPA axis in sheep. PLoS One, 14(1), e0211363.

Verbeek, E., Oliver, M. H., Waas, J. R., McLeay, L. M., Blache, D., & Matthews, L. R. (2012). Reduced cortisol and metabolic responses of thin ewes to an acute cold challenge in mid-pregnancy: implications for animal physiology and welfare. PLoS One, 7(5), e37315.

Vickery, B. H. (1986). Comparison of the potential for therapeutic utilities with gonadotropin-releasing hormone agonists and antagonists. Endocrine reviews, 7(1), 115–124.

Wagner, R., Fieseler, H., Kaiser, M., Müller, H., Mielenz, N., Spilke, J., Gottschalk, J., Einspanier, A., Palme, R., & Rizk, A. (2021). Cortisol concentrations in sheep before, during and after sham foot trimming on a tilt table-the suitability of different matrices. Schweizer Archiv fur Tierheilkunde, 164(11), 753–766.

Walker, D. (1950). The influence of sex upon carcass quality of New Zealand fat lamb. New Zealand Journal of Science and Technology, Section A, 32(1), 30–38.

Weaver, S. J., Hynd, P. I., Ralph, C. R., Edwards, J. H., Burnard, C., Narayan, E., & Tilbrook, A. J. (2021). Chronic elevation of plasma cortisol causes differential expression of predominating glucocorticoid in plasma, saliva, fecal, and wool matrices in sheep. Domestic Animal Endocrinology, 74, 106503.

Wennig, R. (2000). Potential problems with the interpretation of hair analysis results. Forensic science international, 107(1-3), 5–12.

Wester, V. L., van der Wulp, N. R., Koper, J. W., de Rijke, Y. B., & van Rossum, E. F. (2016). Hair cortisol and cortisone are decreased by natural sunlight. Psychoneuroendocrinology, 72, 94–96.

Wojtas, K., Cwynar, P., & Kołacz, R. (2014). Effect of thermal stress on physiological and blood parameters in merino sheep. Journal of Veterinary Research, 58(2), 283–288.

Wright, L., & Perrot, T. (2012). Stress and the developing brain (Vol. 9). Morgan & Claypool Publishers.

Yang, H. Z., Lan, J., Meng, Y. J., Wan, X. J., & Han, D. W. (1998). A preliminary study of steroid reproductive hormones in human hair. The Journal of steroid biochemistry and molecular biology, 67(5-6), 447–450.

Yardimci, M., Sahin, E., Cetingul, I., Bayram, I., Aslan, R., & Sengor, E. (2013). Stress responses to comparative handling procedures in sheep. Animal, 7(1), 143–150.

Yilmaz, O., Demirel, G., Tuzun, A. E., Kara, N. K., & Ekiz, B. (2023). Evaluation of pasture-based vs. concentrate-based lamb production systems regarding growth performance, wool cortisol concentration, meat quality and microbiological properties. Small Ruminant Research, 226, 107026.

Zeinstra, E. C., Vernooij, J., Bentvelzen, M., van der Staay, F. J., & Nordquist, R. E. (2023). Wool cortisol as putative retrospective indicator of stress in ewes during the third trimester of pregnancy, and their newborns: effects of parity and litter size—an exploratory study. Frontiers in Animal Science, 4, 1056726.

Zoratti, A., Corazzin, M., Bodas, R., Domínguez, E., Geß, A., Prandi, A., & Peric, T. (2023). Wool cortisol concentrations trends in the lamb from birth to slaughter. Small Ruminant Research, 224, 106988.

